# Impact of temperature and time on DNA-free Cas9-ribonucleoprotein mediated gene editing in wheat protoplasts and immature embryos

**DOI:** 10.1101/2022.04.05.487229

**Authors:** Snigdha Poddar, Jaclyn Tanaka, Katherine L.D. Running, Gayan K. Kariyawasam, Justin D. Faris, Timothy L. Friesen, Myeong-Je Cho, Jamie H. D. Cate, Brian Staskawicz

## Abstract

The advancement of precision engineering for crop trait improvement is important in the face of rapid population growth, climate change, and disease. To this end, targeted double-stranded break technology using RNA-guided Cas9 has been adopted widely for genome editing in plants. *Agrobacterium* or particle bombardment-based delivery of plasmids encoding Cas9 and guide RNA (gRNA) is common, but requires optimization of expression and often results in random integration of plasmid DNA into the plant genome. Recent advances have described gene editing by the delivery of Cas9 and gRNA as pre-assembled ribonucleoproteins (RNPs) into various plant tissues, but with moderate efficiency in resulting regenerated plants. In this report we describe significant improvements to Cas9-RNP mediated gene editing in wheat. We demonstrate that Cas9-RNP assays in protoplasts are a fast and effective tool for rational selection of optimal gRNAs for gene editing in regenerable immature embryos (IEs), and that high temperature treatment enhances gene editing rates in both tissue types. We also show that Cas9-mediated editing persists for at least 14 days in gold particle bombarded wheat IEs. The regenerated edited wheat plants in this work are recovered at high rates in the absence of exogenous DNA and selection. With this method, we produce knockouts of a set of three homoeologous genes and two pathogenic effector susceptibility genes that result in insensitivity to corresponding necrotrophic effectors produced by *Parastagonospora nodorum*. The establishment of highly efficient, DNA-free gene editing technology holds promise for accelerated trait diversity production in an expansive array of crops.

## Introduction

Amidst a rapidly growing population and threats posed by climate change and disease, there exists a need for the advancement of crop biotechnology to increase the speed and precision of crop varietal development. Cas9 has emerged as a plant gene editing tool of choice for its accuracy and programmability to engineer allelic diversity for beneficial traits to support global food security. Guided by RNA, Cas9 efficiently makes sequence-specific double-stranded breaks in genomic DNA (Jinek et al. 2012). The host’s double-stranded break repair mechanisms are then elicited. Non-homologous end joining (NHEJ), the predominant and often error prone pathway in plants, can lead to insertions or deletions (indels) at the Cas9 cut site upon repair (Puchta 2005). Exploitation of this system allows for targeted knockout of endogenous genes.

Cas9 and guide RNA (gRNA) encoding plasmid DNA systems have been developed and delivered to plant and major crop species including *Arabidopsis* (Li et al. 2013), potato (Wang et al. 2015; Butler et al. 2015), tomato (Brooks et al. 2014; Lor et al. 2014), soybean (Jacobs et al. 2015), maize (Svitashev et al. 2015; Char et al. 2017), barley (Lawrenson et al. 2015; Garcia-Gimenez et al. 2020), rice (Feng et al. 2013), and wheat (Wang et al. 2014) by *Agrobacterium tumefaciens* or particle bombardment. These methods rely on random integration of Cas9-gRNA cassettes into the genome, and optimization of expression for each plant system. As a result, the gene editing process is encumbered by variables such as promoter and terminator choice when cloning constructs and copy number and integration location of transgenes upon transformation. Additionally, gene editing by these methods raise transgenic regulatory concerns. Regulation aside, transgenes can often be segregated away through breeding, but the process is laborious, time consuming, and particularly difficult for plants with complex genomes. Moreover, crops with lengthy generation times or those that are vegetatively propagated, such as cassava and banana, cannot be bred to segregate transgenes. There have been reports in which plant gene editing has been achieved by transient expression of Cas9 and gRNA (Zhang et al. 2016; Hamada et al. 2018), however full experimental control over the fate of transgene integration and tracking has not been achieved. For these reasons, there is a clear need for advances in DNA-free genome editing technology.

The direct delivery of preassembled Cas9-gRNA ribonucleoproteins (RNPs) is one such technology and has been demonstrated in various plant protoplast systems to induce targeted mutations (Woo et al. 2015; Malnoy et al. 2016; Poddar et al. 2020; Brandt et al. 2020; Sant’Ana et al. 2020). Some have produced edited plants arising from the transfected single cells. However regeneration of wheat and other crop plant protoplasts is not feasible with current methods.

Cas9-RNP based editing of maize (Svitashev et al. 2016), rice (Banakar et al. 2019), and wheat (Liang et al. 2017) regenerable embryos by biolistics has also been reported. Gold particles coated with Cas9-RNPs are bombarded with high pressure into immature embryos (IEs) that are ultimately regenerated into plants through tissue culture. Co-delivery of DNA vectors with selective markers or helper genes along with Cas9-RNPs have been utilized to improve editing efficiency (Svitashev et al. 2016; Banakar et al. 2019). In the absence of selection, however, editing rates have generally been low.

The use of Cas9-RNPs to generate edited plants provides unique benefits. Because the gene editing reagents are delivered as pre-assembled complexes, researchers do not need to optimize DNA vectors, the host plant tissue does not bear the burden of transcribing or translating Cas9 or gRNA, and breeding for segregation is unnecessary due to the absence of transgenes. Additionally, the Cas9-RNPs, which exist in a finite amount in the target tissue, are ultimately degraded by endogenous proteases and nucleases. However, there remains room to improve the editing pipeline and increase efficiency.

Low rates of Cas9 mediated editing in plant tissue may indicate that the endonuclease is not reaching its full potential due to suboptimal environmental conditions. For example, studies across organisms including *Arabidopsis*, citrus (LeBlanc et al. 2018), and wheat (Milner et al. 2020) have shown that Cas9 generates more targeted indels at elevated temperatures.

Here, we present advances in Cas9-RNP based gene editing in the global food crop, wheat (*Triticum aestivum*). To determine if temperature can be harnessed to enhance Cas9-RNP mediated editing, we explore the effects of heat treatment on transfected wheat protoplasts and IEs. We examine the relationship of editing efficiency between non-regenerable protoplasts and regenerable IEs and monitor the rate of editing over time. We demonstrate that treatment at elevated temperatures increases gene editing efficiency in both tissue systems and find that the RNP transfection technique of gold particle bombardment results in sustained editing of tissue at least 14 days after bombardment. We also find that editing rates in protoplasts correlate linearly with editing rates in IEs. Therefore, rapid *in vivo* protoplast assays can be instituted as a standard gene editing pipeline step to select the most effective gRNAs for IE gene editing and regeneration. Lastly, we regenerate wheat plants edited via Cas9-RNP biolistic transfection. As a proof of method, we simultaneously target three wheat homoeologous orthologs of a rice gene, *Pi21*(Os04g0401000), and successfully generate lines with knockouts in all copies. We also target wheat genes *Tsn1* and *Snn5*, producing lines that are insensitive to the *Parastagonospora nodorum* pathogenic effectors SnToxA and SnTox5 and establish DNA and selection-free Cas9-RNP mediated editing as an efficient and feasible technique for generating targeted gene knockouts in wheat.

## Results

### Cas9-RNP transfection and the effect of temperature in wheat protoplast gene editing

We first quantified cell viability after heat treatment of non-transfected protoplasts to determine the feasibility of testing higher temperatures for wheat protoplast gene editing. Protoplasts were isolated from partially etiolated wheat seedlings and incubated at 25°C, 30°C, or 37°C for 16 hours followed by 25°C for 8 hours. During the 24-hour period, the protoplasts were monitored for viability every 8 hours using Evans blue staining and microscopy. Viability of protoplasts treated at 37°C decreased markedly compared to those treated at 25°C and 30°C (Figure S1) and suffered from media evaporation. It was therefore concluded that the protoplast gene editing pipeline was not amenable to a 37°C heat treatment.

Five single guide RNAs (sgRNAs), Pi21gD, Tsn1g2, Tsn1g3, Snn5g1, and Snn5g2 were selected and commercially synthesized for this study. To assess the efficacy of the sgRNAs *in vivo*, and to determine the effect of temperature on wheat protoplast gene editing, Cas9-RNPs were assembled and transfected into wheat mesophyll protoplasts. Purified Cas9 with a C-terminal double nuclear-localization tag was complexed with sgRNA. The resulting sgRNA-Cas9 RNPs were transfected into wheat protoplasts using polyethylene glycol (PEG). Transfected protoplasts were treated at 25°C or 30°C and harvested for genotypic analysis after 24 hours. Editing rates at the target loci were determined by amplicon next-generation sequencing (NGS). With incubation at 25°C and 30°C, average editing rates ranged from 2.5-50% and 5.8-62% respectively. Despite this variability between different sgRNA-Cas9 RNPs, editing efficiency was consistently higher in protoplasts treated at 30°C compared to 25°C for any given sgRNA (Figure 1), suggesting that a higher temperature treatment is advantageous to RNP-mediated gene editing in wheat protoplasts.

**Figure 1.**
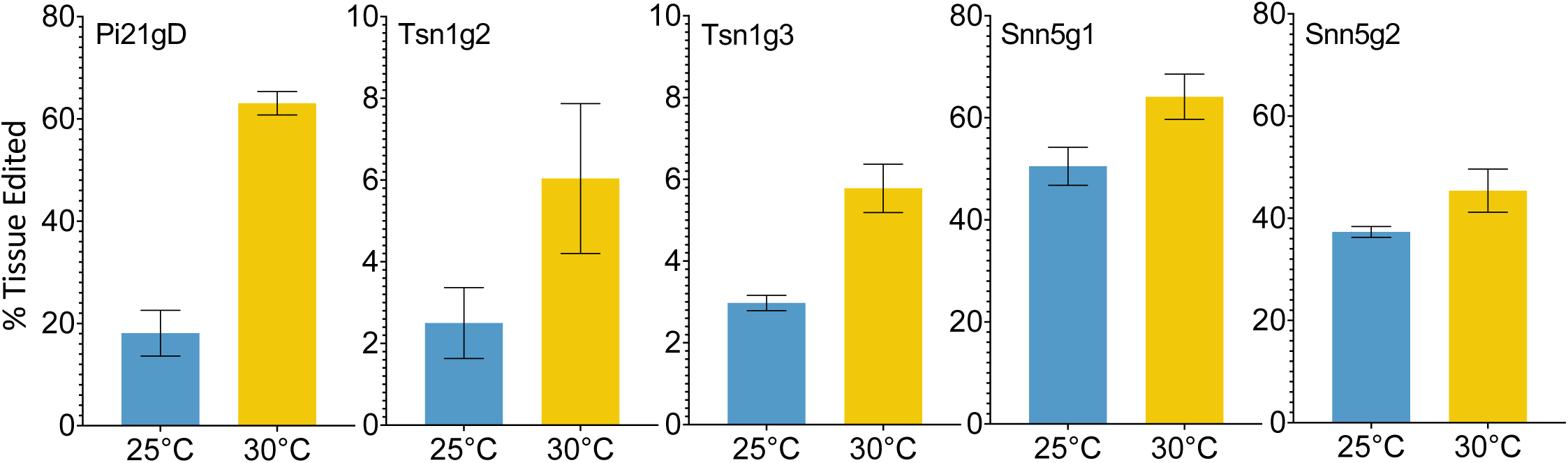
Targeted editing efficiency of gRNA-Cas9 ribonucleoproteins with different temperature treatments in wheat protoplasts. Five gRNAs, Pi21gD, Tsn1g2, Tsn1g3, Snn5g1, and Snn5g2 were tested and transfected independently into protoplasts. N=3. Error bars indicate SEM.

### Biolistic Cas9-RNP delivery and the effect of temperature in wheat immature embryo gene editing

To determine if a high temperature treatment similarly improves Cas9-RNP based editing in wheat IEs as it does in protoplasts, RNPs were transfected into IEs by particle bombardment. The experimental pipeline is summarized in Figure 2a. Single guide RNA and Cas9 were complexed *in vitro*, adsorbed onto 0.6 μm gold particles, and biolistically delivered with a helium-pressured particle gun. For each sgRNA and temperature being tested, 30 IEs were bombarded and incubated at 26°C, 30°C, or 37°C for 16 hours. They were then maintained at 26°C on callus-induction media before inducing regeneration at around 63 days post-bombardment (dpb). Plasmid DNA was not co-delivered with any of the Cas9-RNPs, and callus induction and regeneration were performed under selection-free conditions. From each set of 30 RNP-transfected embryos, ten were randomly harvested and pooled for genomic analysis at 14 dpb and again at 48 dpb. The remaining ten embryos were kept for regeneration into M_0_ plants. All independent shoots were isolated and treated as individual M_0_ plants. Plants were transplanted from tissue culture media to soil approximately 100 dpb. Each resulting M_0_ plant was independently genotyped, and the percent tissue edited rate was calculated as the percentage of mutant alleles among total alleles in the M_0_ plant pool. The percentage of plants edited was also calculated as a percentage of the number of plants with any edit among the number of total M_0_ plants regenerated. All genomic analysis was done by amplicon NGS.

**Figure 2.**
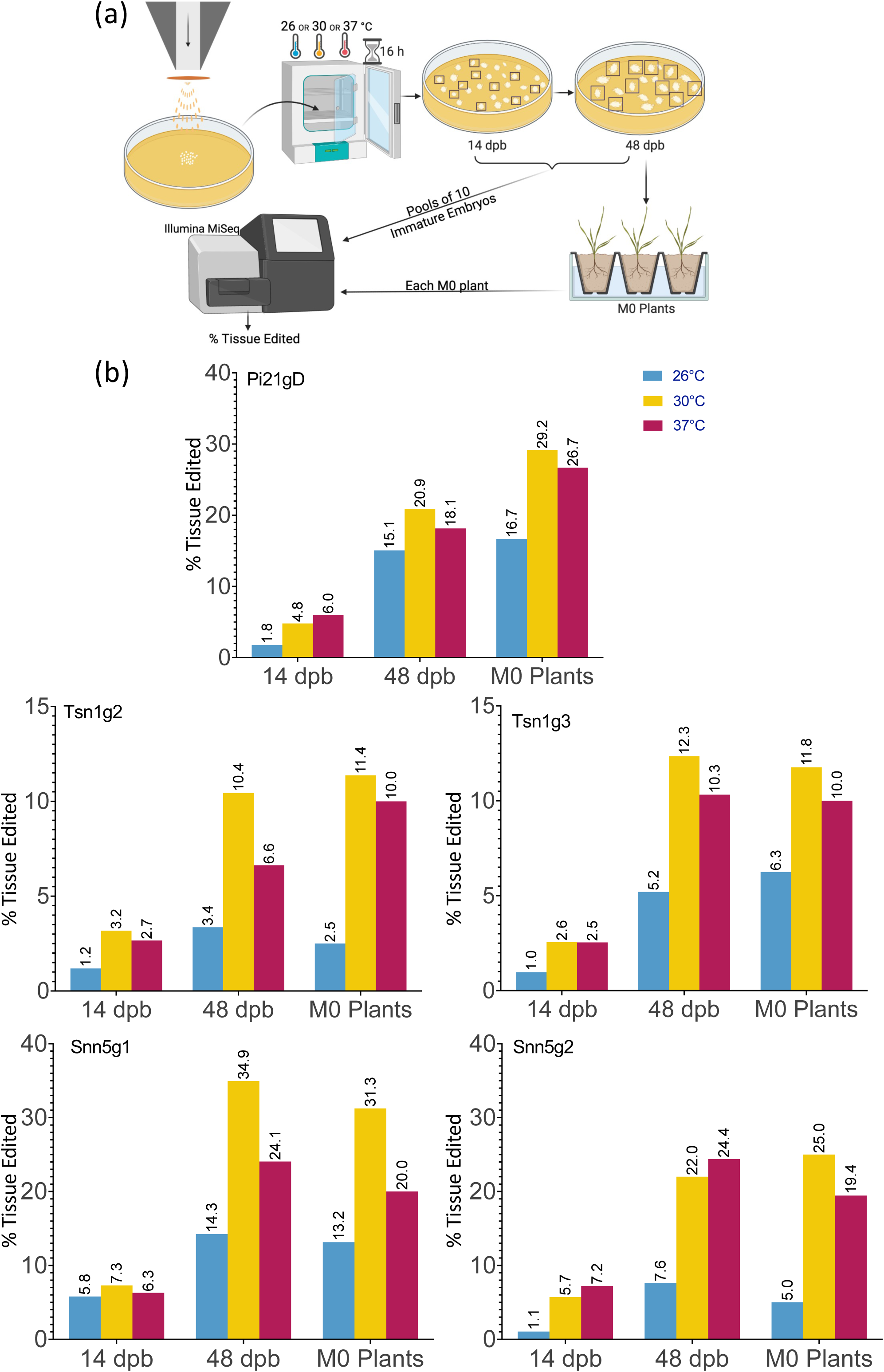
Cas9-RNP particle bombardment and temperature treatment of wheat immature embryos (IEs). (a) A schematic of the particle bombardment and editing efficiency assay pipeline. (b) Targeted editing efficiency of gRNA-Cas9 ribonucleoproteins with different temperature treatments in IEs across time points. Five gRNAs, Pi21gD, Tsn1g2, Tsn1g3, Snn5g1, and Snn5g2 were bombarded independently into IEs. Tissue pools at 14 dpb and 48 dpb consisted of 10 randomly chosen initially bombarded IEs. Editing efficiency for M_0_ plants is based on aggregate data from all independently genotyped M_0_ plants that emerged from 10 randomly chosen initially bombarded IEs. Percent tissue edited is defined as the percentage of tissue with insertions or deletions within 2 bp of the target cleavage site out of the total tissue pool.

Elevated temperature treatment of both 30°C and 37°C led to higher percentages of edited tissue compared to 26°C for all five sgRNA-Cas9 RNPs across all timepoints (Figure 2b). Tissue editing rates were higher at 48 dpb than at 14 dpb and editing rates in the M_0_ regenerant tissue pool were comparable to those at 48 dpb. From the ten embryos per treatment allowed to regenerate, 10-40 M_0_ plants were produced. Plants with wild type, heterozygous, biallelic, and homozygous mutations at the target loci were obtained. Editing efficiency in the M_0_ regenerants is summarized in Table 1 and genotypes of each individual edited M_0_ regenerant are described in Table S3.

**Table 1.**
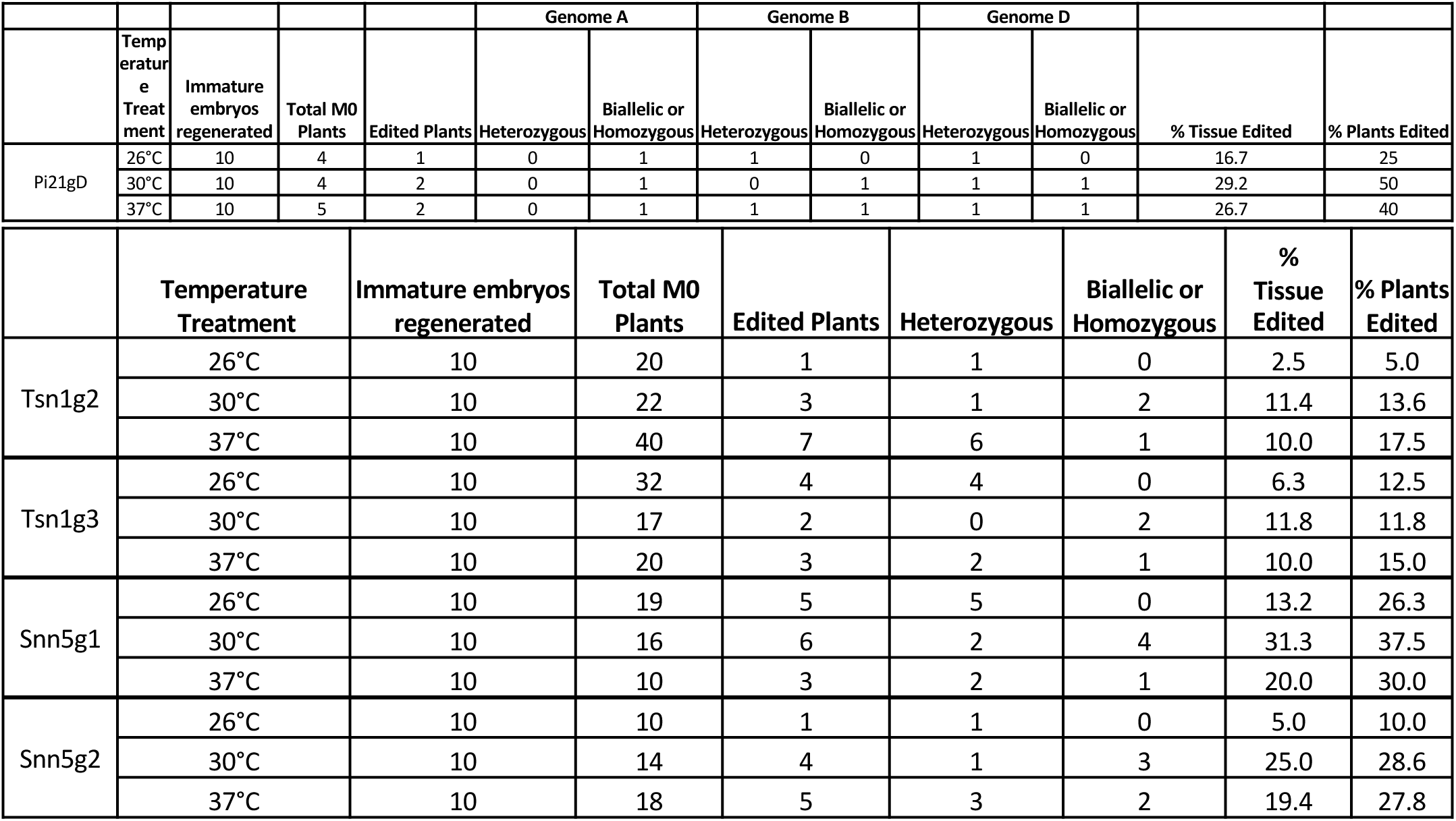
Summary of editing outcomes in *Pi21, Tsn1*, and *Snn5* targeted M_0_ plants. Data for *Pi21* is broken down by subgenome. *Tsn1* and *Snn5* are only present on subgenome B. “% Tissue Edited” indicates the percentage of edited alleles among the total alleles analyzed from the M_0_ pools. “% Plants Edited” indicates the percentage of plants with any level of editing among the total plants analyzed from the M_0_ pools.

### Cas9-RNP mediated editing is sustained over time

Notably, gene editing rates were more than doubled, regardless of temperature treatment, in tissue assayed at 48 dpb compared to 14 dpb (Figure 2b). To further investigate the difference in editing rates over time, the number of unique mutant alleles was determined at the 14 and 48 dpb timepoints. With minimal exception, there were more unique mutant alleles at 48 dpb compared to 14 dpb (Figure 3a, Figure S2).

**Figure 3.**
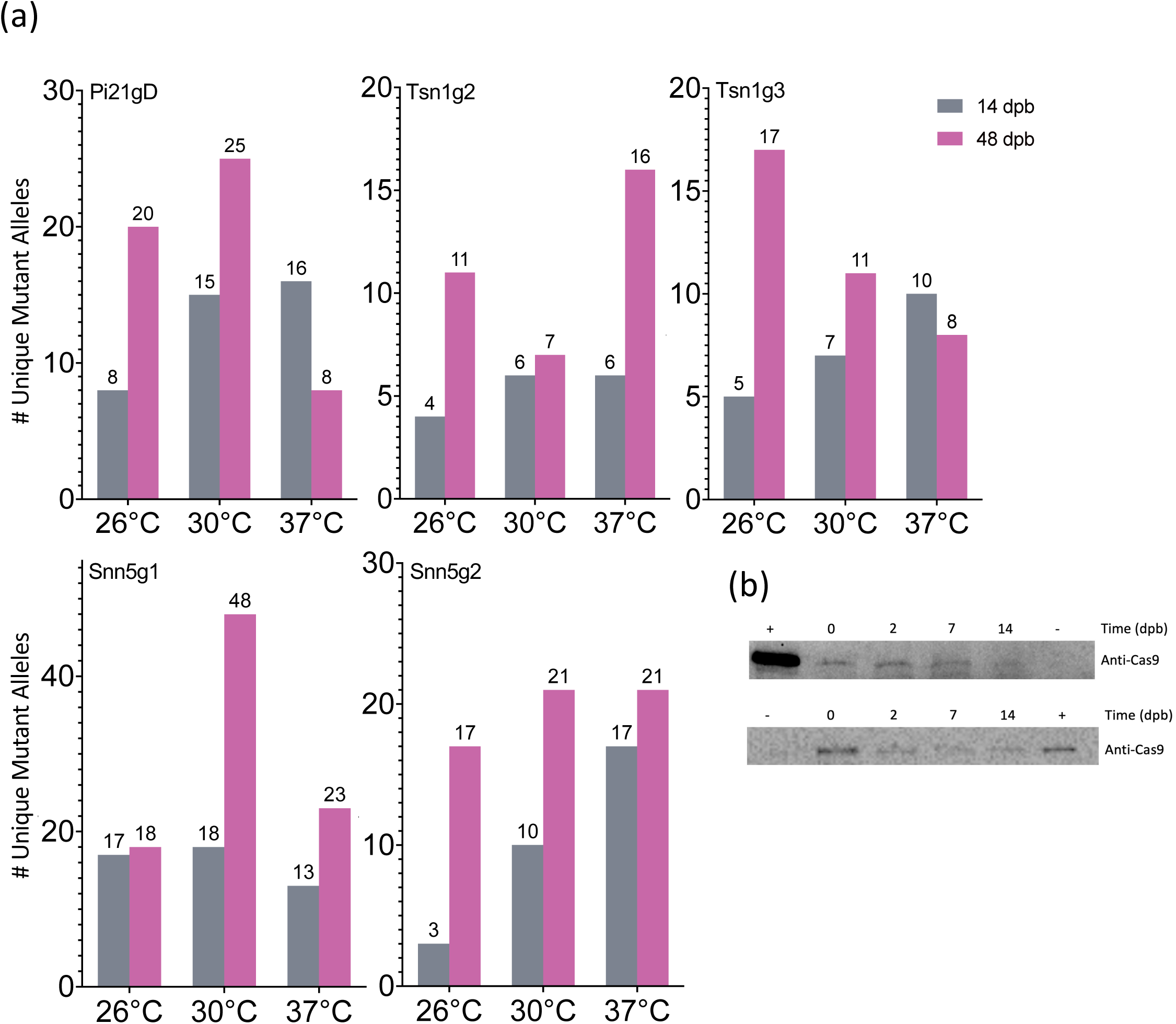
Cas9-RNP mediated editing in gold particle bombarded immature embryos IEs is sustained over time. (a) Quantification of the number of unique mutant alleles detected via deep sequencing. (b) Western blot detection of Cas9 in 10-IE bombarded samples taken 0, 2, 7, and 14 dpb with anti-Cas9 antibody. The top and bottom blot represent 2 independent sets of 10 IEs. + = 9 ng (top) and 3 ng (bottom) Cas9; - = IEs that were not bombarded with Cas9-RNP; loading volume of 25 μl (top) and 40 μl (bottom) total soluble protein extract per IE sample.

An additional 50 IEs were bombarded with Snn5g1-Cas9 RNP to determine the length of time that Cas9 remains present in biolistically transfected tissue. Western blot analysis was performed with 10-embryo tissue samples taken 0, 2, 7, and 14 dpb. Given the finite amount of Cas9 protein delivered by RNP bombardment and rapid cell division and growth in each IE over time, we normalized the experiment by volume extracted from total tissue originating from ten IEs at any given timepoint, rather than total protein extracted. Cas9 was detected in tissue from all four timepoints with decreasing band intensity over time (Figure 3b). Cas9 was not detected in embryos that were not subjected to bombardment of Cas9-RNPs. Due to the large mass of tissue from exponential growth of callus from IEs, it was not feasible to extract protein from and perform Western blot analysis on ten-embryo 48 dpb samples. Taken together, these results suggest that Cas9 mediated editing activity is sustained over the course of at least 14 days after biolistic delivery of Cas9-RNPs into immature wheat embryos. When using this method, the degradation of Cas9 protein in the target tissue is not as rapid as previously hypothesized (Kim et al. 2014), and evaluation of editing efficiency should occur 14 to 48 dpb for increased accuracy.

### Relative editing rates in protoplasts correlate linearly with editing rates in M_0_ regenerants from bombarded immature embryos

The different sgRNA-Cas9 RNPs used in this study conferred different levels of efficacy in both PEG transfected protoplasts and biolistically transfected embryos. To determine whether the editing rates in the two tissue systems correlated with one another, each sgRNA-Cas9 RNP’s average editing efficiency in 30°C treated protoplasts was plotted against its editing efficiency in 48 dpb 30°C treated bombarded IEs as well as the M_0_ 30°C treated regenerant tissue pool. A linear regression model was applied to the data, revealing a positive linear correlation with *R*^2^=0.744 and *R*^2^= 0.994, respectively (Figure 4). Though a survey of a greater number of sgRNAs would strengthen this association, the present data suggest that editing efficiency in protoplasts can be predictive of editing efficiency in IEs. Given the positive correlation between RNP-mediated editing rates in protoplasts and in biolistically transfected IEs, it can be beneficial to first rapidly score the efficiency of various gRNA candidates in protoplasts to optimize for the highest rate of edited regenerant tissue.

**Figure 4.**
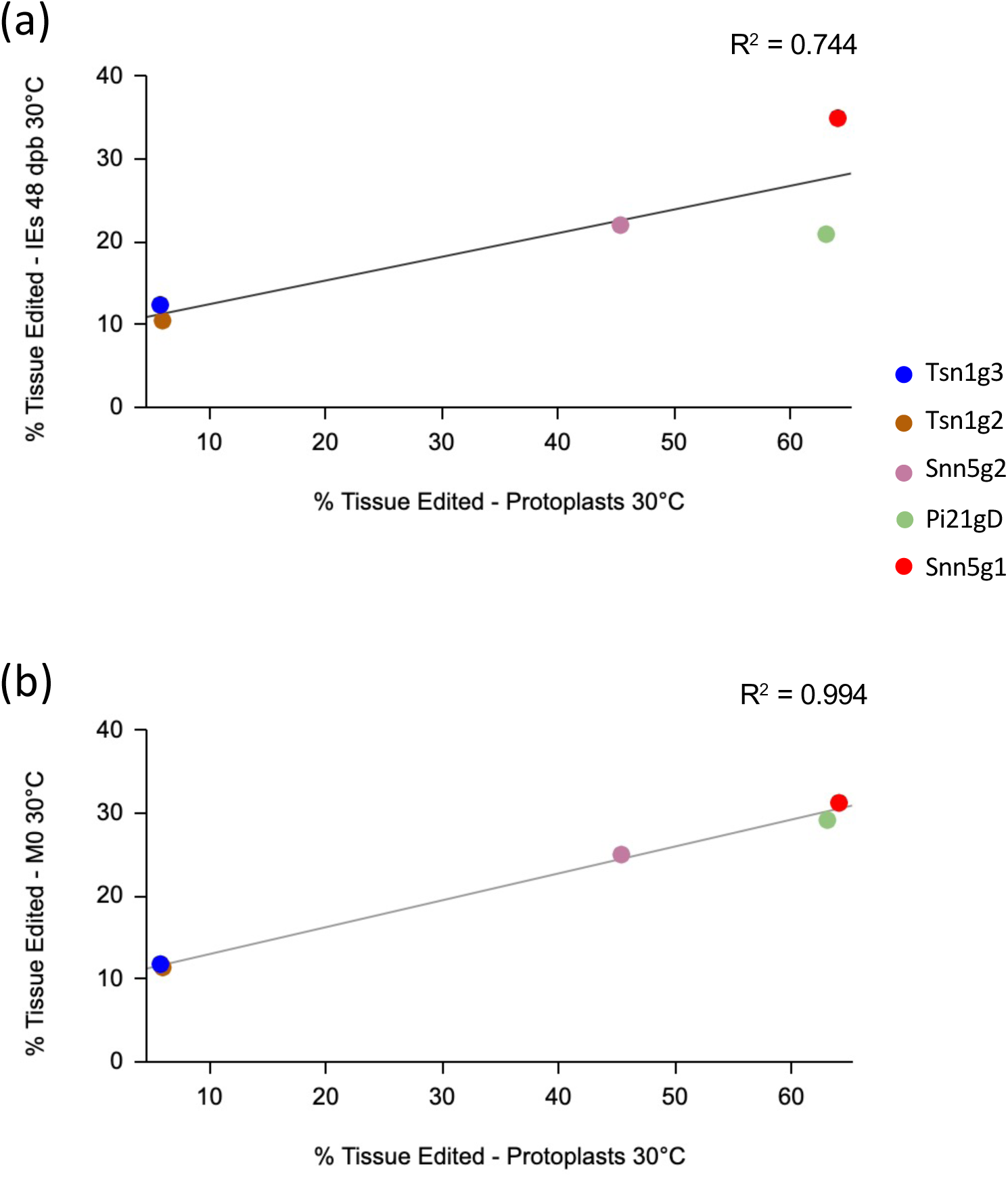
Correlation plot between targeted editing efficiency of gRNA-Cas9 RNPs in protoplasts and immature embryos (IEs) at **(a)** 48 dpb and in **(b)** M_0_ plants treated at 30°C.

### Cas9-RNP mediated knockout of *Parastagonospora nodorum* necrotrophic effector sensitivity genes

The wheat genes *Tsn1* and *Snn5* recognize necrotrophic effectors produced by *Parastagonospora nodorum*, and each exist as single copy genes on the B genome of allohexaploid wheat. In this study, 20 M_0_ *Tsn1* edited plants were produced from 30 transfected embryos maintained for regeneration. Of those, 14 had heterozygous mutations and 6 had biallelic or homozygous mutations. Fully expanded secondary leaves of a subset of M_0_ *Tsn1* edited plants, M_0_ *Tsn1* WT plants, and Fielder grown from seed were infiltrated with SnToxA expressed in *Pichia pastoris*. After 72 hours, M_0_ heterozygotes, M_0_ WT, and Fielder plants had necrotic lesions extending from the site of infiltration. Meanwhile, M_0_ plants with biallelic or homozygous mutations exhibited no necrosis (Figure 5).

**Figure 5.**
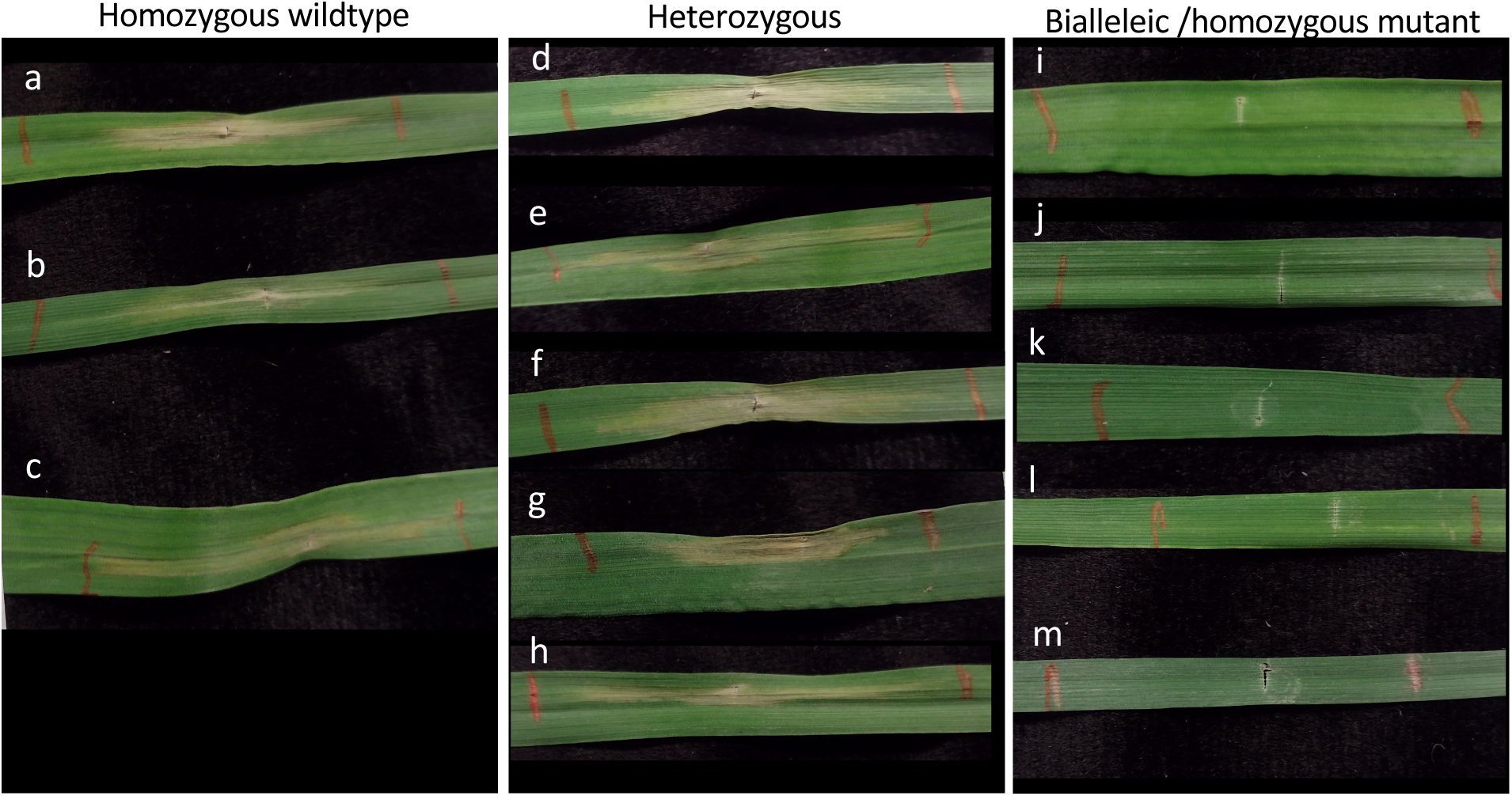
SnToxA assay in *Tsn1* targeted M_0_ regenerants. (a) Fielder control grown from seed. (b-m) independent M_0_ regenerants with (b, c) homozygous wildtype; (d-h) heterozygous (d) -2; (e) -5; (f) -31; (g) -1; (h) +1; and (i-m) biallelic or homozygous mutant (i) -2, -5; (j) -2, -2, (k) -1, -1; (l) -2, -2; (m) -1, -1 genotypes. Mutation notation is as follows: a positive number, +, indicates the number of bases inserted, a negative number, -, indicates the number of bases deleted.

Similarly, a total of 24 M_0_ *Snn5* edited plants were produced from 30 transfected embryos maintained for regeneration. Of those, 14 had heterozygous mutations and ten had biallelic or homozygous mutations. Fully expanded secondary leaves of a subset of M_0_ *Snn5* edited plants, M_0_ *Snn5* WT plants, and Fielder grown from seed were infiltrated with SnTox5 containing culture filtrates. After 72 hours, M_0_ heterozygotes with in-frame deletions, M_0_ WT, and Fielder plants exhibited necrotic lesions. Results for M_0_ heterozygotes, however, displayed a mixture of phenotypes ranging from sensitive to insensitive. Two heterozygous plants with an in-frame deletion on one allele appeared insensitive to SnTox5. Notably, all plants with biallelic or homozygous mutations leading to premature termination were insensitive to SnTox5 (Figure 6).

**Figure 6.**
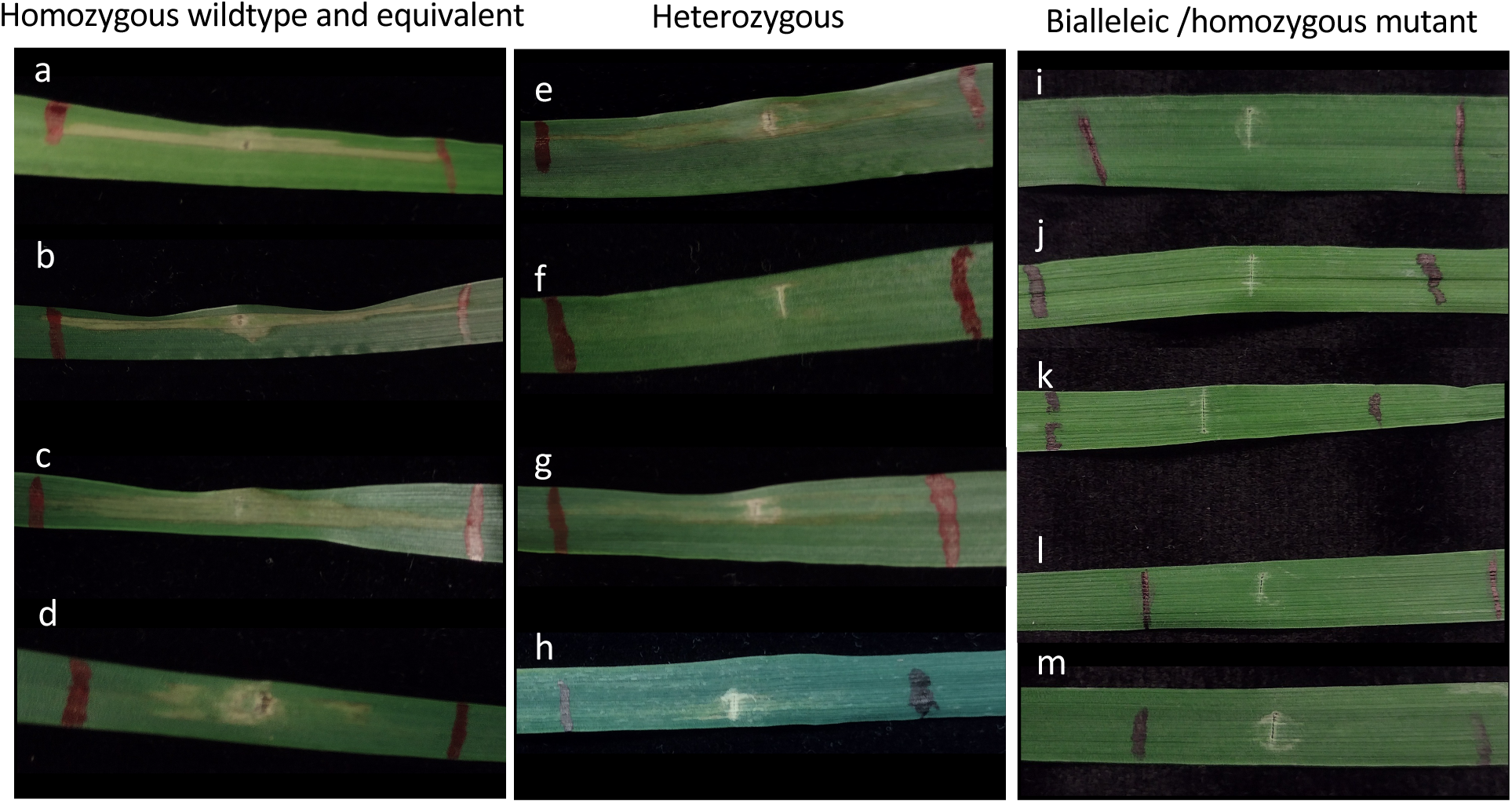
SnTox5 assay in *Snn5* targeted M_0_ regenerants. (a) Fielder control grown from seed. (b-m) independent M_0_ regenerants with (b) homozygous wildtype; (c-d) heterozygous in-frame mutant: (c) -3; (d) -6; (e-h) heterozygous mutant: (e) -5; (f) +20; (g) +2-1; (h) -4; (i-m) biallelic or homozygous mutant: (i) -11, -4; (j) -8, -2; (k) -10, -10; (l) +1, -2; (m) -5, -1 genotypes. Mutation notation is as follows: a positive number, +, indicates the number of bases inserted, a negative number, -, indicates the number of bases deleted.

These results demonstrate that loss-of-function mutations can be introduced to both copies of a gene within the M_0_ generation, leading to insensitivity to agronomically relevant necrotrophic fungal effectors. M_0_ heterozygotes and biallelic plants can be self-fertilized to establish lines with homozygous deleterious mutations in the susceptibility genes. The biolistic method with 30°C or 37°C heat treatment is highly efficient, and edited plants can be identified from a small number of regenerants without the use of selection in tissue culture.

## Discussion

CRISPR-based RNPs have been used for editing in various plant species and tissue types (Zhang et al. 2021). In this work, we improve upon DNA-free Cas9-RNP technology for genome editing in wheat. We establish heat treatment as a parameter to increase the rate of editing *in vivo*, show that particle bombardment-based editing is sustained over more than 14 days, and demonstrate that results from protoplast assays can be utilized as a proxy for predicting editing rates in regenerable tissue and as a tool to rank gRNA efficacy. By delivering gene editing reagents as protein-RNA complexes, several complications associated with *Agrobacterium tumefaciens* and biolistic DNA vector delivery are avoided.

Cas9 from *Streptococcus pyogenes*, a bacterium that grows optimally at 37°C (Zhou & Li 2015), has been shown to exhibit increased cleavage activity at 37° compared to 22°C *in vitro* (LeBlanc et al. 2018). Plant protoplast and IE transfections and regeneration are typically performed at ambient temperatures (25°C and 26°C respectively). Although modulation of temperature has not been previously performed in protoplast gene editing experiments, an increase in temperature for DNA-based plant gene editing studies have resulted in higher targeted mutation frequencies (LeBlanc et al. 2018; Malzahn et al. 2019). The application of temperature treatment to increase Cas9-RNP mediated editing efficiency in any plant tissue system has not previously been demonstrated. Here, we found that 16 hours of exposure of Cas9-RNP transfected protoplasts to 30°C markedly increased indel formation at the Cas9 cut site (Figure 1). Similarly, 16 hours of exposure of Cas9-RNP bombarded IEs to 30°C or 37°C resulted in increased targeted indel formation. In IEs assayed at 48 dpb we achieved editing rates of 10.4-34.9% with 30°C treatment, 6.63-24.39% with 37°C treatment, and just 3.36-14.25% with standard 26°C incubation (Figure 2b). Interestingly, the benefit of increased temperature treatment was consistent between the two target tissues and across the five different target sites tested. In our work, there were no discernable defects in regenerability for IEs treated at a higher temperature compared to the standard 26°C. We detected no positive or negative correlation between temperature treatment and the number of M_0_ plants recovered.

Two reports have described the biolistic delivery of Cas9-RNPs into wheat and maize embryo cells in the absence of DNA and selection (Liang et al. 2017; Svitashev et al. 2016). Both achieved moderate targeted mutagenesis frequencies in the regenerated plants. We noted that the studies each assayed for editing efficiency in the IEs 2 dpb and universally achieved <1% targeted editing. In contrast, the editing efficiencies in regenerated plant tissue were substantially higher, ranging from 1.3-4.7% (Liang et al. 2017) and 2.4-9.7% (Svitashev et al. 2016). To investigate this discrepancy between timepoints, we monitored editing efficiency at 14 dpb, 48 dpb, and in the M_0_ regenerants in our study. Irrespective of temperature treatment or gRNA sequence, editing frequencies at 48 dpb were considerably higher than at 14 dpb (Figure 2b). Percentage of tissue edited in the M_0_ plant pool was comparable to that at 48 dpb. The observed difference in editing efficiency between earlier timepoints and regenerated M_0_ plants was consistent with previous reports (Liang et al. 2017; Svitashev et al. 2016).

In mammalian cells, Cas9 was shown to be undetectable 48-72 hours after Cas9-RNP transfection by nucleofection (Kim et al. 2014). For this reason, it has been thought that enzymatic degradation of Cas9-RNPs *in vivo* is rapid and that editing must occur within the first few days of transfection. In the present study, if Cas9-RNPs were fully degraded from the tissue prior to the 14 dpb timepoint, all gene editing would have had to occur before 14 dpb. Consequently, approximately the same number of unique alleles would have been expected to be detected at both 14 dpb and 48 dpb if proliferation of edited and unedited cells occurs at the same rate. On the contrary, consistently higher rates of mutagenesis as well as a greater number of unique alleles at the later timepoints were observed at 48 dpb (Figure 2b, Figure 3a, Figure S2), suggesting that Cas9 may somehow be stabilized for at least 14 days and gradually released within the wheat IEs after biolistic delivery for sustained editing over time. As further evidence in support of this hypothesis, Cas9 protein was detected in 10-embryo tissue samples taken 2, 7, and 14 dpb (Figure 3b). Taken together, these results indicate that Cas9 is maintained in tissue at least 14 dpb and facilitates sustained and gradual editing of tissue over time when delivered as Cas9-RNP via gold particle bombardment. Further biochemical exploration is necessary to understand the mechanism of this Cas9 stabilization and persistent editing.

Numerous plant protoplast systems have been used for targeted mutagenesis using Cas9-RNPs (Woo et al. 2015; Malnoy et al. 2016; Shan et al. 2019; Poddar et al. 2020; Brandt et al. 2020; Sant’Ana et al. 2020; Yu et al. 2021). Although the method is useful for producing Cas9-RNP edited plants for protoplasts that are amenable to regeneration, most crop plants cannot easily be regenerated in this manner. Though wheat protoplasts are recalcitrant to regeneration through existing methodology, protoplasts in the current study prove to be a beneficial screening system. Cas9-RNP mediated editing rates in protoplasts correlated linearly with editing rates in IEs. Because biolistic Cas9-RNP transfection of IEs requires significant time, energy, resources, and commitment, a means for rational selection of gRNA sequences for optimal editing efficiency is preferred. It is noteworthy that there were major differences in mutation rates for the 5 gRNAs used in this study. Unfortunately, existing predictive software to select gRNAs often do not translate upon experimentation. Therefore, when attempting to select the best gRNA to produce the highest rate of stable editing in regenerable IEs, transient protoplast assays can serve as a rapid pipeline to rank gRNAs and forecast editing rates in Cas9-RNP bombarded regenerable tissue.

The calculation of editing efficiency in M_0_ regenerants has the potential to be confusing. To be explicit in our analysis, we present editing rates of regenerants in two ways. The percentage of total edited alleles in the M_0_ regenerant pool is indicated as “% Tissue edited”, while “% Plants edited” is the percentage of total edited plants among all the M_0_ plants (Figure 2b, Figure 4, Table 1). The former is meant to compare overall editing efficiency more fairly across tissue types and timepoints, taking biallelism, homozygosity, and heterozygosity of regenerated plants into consideration. The latter value is more relevant for evaluating the method’s ability to produce individual plants with gene edits.

The gene *Pi21* was first characterized in rice (*Oryza sativa*) as a negative regulator of resistance for blast disease (Fukuoka et al. 2009). We identified putative orthologs in wheat that consisted of three homoeologous genes. The functionality of wheat *Pi21* has not been formally assessed but may potentially play a role in disease susceptibility. Wheat *Pi21* was selected as a target to demonstrate the DNA-free Cas9-RNP gene editing method in a gene present in all three diploid subgenomes (AABBDD). Pi21gD was designed to simultaneously target all six alleles. Despite the genetic complexity, we were able to regenerate plants with biallelic or homozygous mutations across all three subgenomes for a full variety of genotypes including two with biallelic or homozygous triple mutant edits within the M_0_ generation (Table S3).

The wheat genes *Tsn1* and *Snn5* recognize the *Parastagonospora nodorum* pathogenic effectors SnToxA and SnTox5, respectively (Faris et al. 2010; Kariyawasam et al. 2021). *Tsn1* is a gene with resistance gene-like features including protein kinase, nucleotide binding, and leucine-rich repeats, and the ToxA necrotrophic effector is produced by at least three economically important fungal pathogens of wheat (Friesen & Faris 2021). *Snn5* belongs to a different class and contains protein kinase and major sperm protein domains (details regarding the cloning and characterization of *Snn5* will be published in the future; K.L.D. Running and J.D. Faris, personal communication), but like *Tsn1*, it functions as a target for a necrotrophic effector leading to disease susceptibility (Kariyawasam et al. 2021). Therefore, *Tsn1* and *Snn5* are practical targets for disruption via DNA-free gene editing. Using DNA-free biolistic delivery of Cas9-RNPs, we successfully generated plants with heterozygous, biallelic, and homozygous mutations within the M_0_ generation from a mere ten IEs per treatment. Biallelic and homozygous mutants of *Tsn1* and *Snn5* were demonstrated to be insensitive to SnToxA and SnTox5, respectively. Due to the high rate of editing, particularly using Snn5g1 and Snn5g2 with 30°C and 37°C heat treatment, screening of M_0_ plants for edits was fully feasible. Contrary to previous reports, a selection scheme can reasonably be foregone with Cas9-RNP mediated editing so long as gRNAs are pre-tested in protoplasts and deemed to be highly effective.

In summary, heat treatment enhancement of Cas9-RNP mediated wheat editing combined with a protoplast-based approach to select optimal gRNAs, and findings that editing is sustained for more than 2 weeks advances this DNA and selection-free gene editing approach in crops. Given the persistence of Cas9 in bombarded tissue, additional work with increased length or punctuated exposure to heat, beyond 16 hours, throughout callus induction may further augment the benefit of heat treatment. The success of this method in targeting single loci warrants exploration of furthering the technique to multiplexing. In addition to knocking out genes, editing via Cas9-RNPs can conceivably be applied to generating allelic series by targeting non-coding genomic regions such as promoters (Rodríguez-Leal et al. 2017). The presented advancement to this technology can be applied to numerous crops that are amenable to particle bombardment and encourages the establishment of tissue culture and regeneration protocols in crop species that are vegetatively propagated.

## Materials and Methods

### Plant material

The allohexaploid wheat (*Triticum aestivum* L., 2*n* = 6*x* = 42, AABBDD genomes) cultivar Fielder was used for this study.

### Cas9-gRNA RNP assembly

Cas9 protein with a C-terminal double nuclear-localization tag (QB3 Macrolab, University of California, Berkeley) and sgRNAs with modifications of 2’-O-Methyl at 3 first and last bases, and 3’ phosphorothioate bonds between first 3 and last 2 bases (Synthego, Menlo Park, CA) were complexed *in vitro* to form Cas9-gRNA RNPs.

For each protoplast transfection, a 25 μl reaction was assembled. Thoroughly mixed were 10 μg sgRNA, 2.5 μl 10X NEBuffer 3.1 (New England Biolabs, Ipswich, MA), and nuclease-free water. Then, in a drop-wise manner, 10 μg Cas9 was added slowly with constant mixing, followed by 20 min incubation at 37°C.

For each IE biolistic transfection, a 40 ul reaction was assembled. Thoroughly mixed were 6.4 μg sgRNA, 4 μl 10X NEBuffer 3.1, and nuclease-free water. Then, in a drop-wise manner, 12.8 μg Cas9 was added slowly with constant mixing, followed by 20 min incubation at 37°C.

The resultant RNP mixtures were stored on ice until transfection.

### Protoplast isolation and transfection

Partially etiolated seedlings were used as donor tissue for protoplast isolation. Seeds were surface sterilized in 20% (v/v) bleach and rinsed in sterile water. Seedlings were grown under sterile conditions on wet filter paper in the dark for 12-14 days at 25°C with exposure to ambient light for 6 hours every 5 days. Wheat protoplasts were isolated from the donor tissue using a previously described method (Shan et al. 2014). For each transfection 25 μl of Cas9-gRNA RNP mixture, as defined above, were added to 5 × 10^5^ protoplasts. PEG-meditated transfection was performed as described in the literature (Shan et al. 2014). Protoplasts were harvested 24 hours post-transfection for analysis.

### Gold particle preparation for bombardment

Cas9-RNPs were precipitated onto 0.6 μm gold particles (#1652262, Bio-Rad, Hercules, CA) using the cationic lipid polymer TransIT-2020 (Mirus, Madison, WI) as previously described (Svitashev et al. 2016), with modifications. Briefly, for each 30-IE transfection, 40 μl Cas9-RNP mixture, as described above, was mixed gently with 20 μl sterile gold particles (10 μg μl^-1^ water suspension) and 1 μl TransIT-2020 and incubated on ice for 20 min. The Cas9-RNP coated gold particles were pelleted in a mini microcentrifuge at 2,000*g* for 30 s. The supernatant was removed, and the gold particles were resuspended in 20 μl of sterile water by brief sonication. The coated gold particles were immediately applied to 2 macrocarriers (10 μl each) by spotting numerous small drops and allowed to air dry in a laminar flow hood. For a single transfection, each 30-IE set was bombarded twice using the 2 prepared macrocarriers.

### Immature embryo bombardment and regeneration

Plants were grown at 24°C, 16-hour days and 15°C, 8-hour nights under light intensity of 130 μmol m^-2^s^-1^. Immature seeds containing IEs, sized 1.7-2.2 mm were harvested from wheat spikes 10-13 days after flowering, surface sterilized in 20% (v/v) bleach with one drop of Tween 20 and triple rinsed with sterile water, followed by extraction of the IEs. The IEs were placed on DBC3 media (Cho et al. 1998), scutellum side up and incubated overnight at 26°C prior to biolistic transfection. Four hours prior to bombardment, IEs were placed on 55 mm filter paper in the center of DBC3 osmoticum media containing 0.2 M mannitol and 0.2 M sorbitol (Cho et al. 2000). Using two prepared microcarriers holding Cas9-RNP coated gold microparticles, IEs were shot twice using the PDS-100/He gene gun (Bio-Rad, Hercules, CA) with rupture pressure of 1100 psi. The bombarded IEs were transferred from the filter paper directly to the media below and incubated at 26°C, 30°C, or 37°C for 16 hours. IEs were transferred to standard DBC3 media in dim light (10 μmol m^-2^ s^-1^) at 26°C for 9 weeks with subculturing as needed. Callus tissue originating from each IE was transferred to DBC6 media for regeneration (Cho et al. 2015). Resultant plantlets were transferred to rooting media and incubated in high light (90 μmol m^-2^ s^-1^) at 26°C and grown to 4-6 inches before being transplanted to soil.

### Amplicon next generation sequencing analysis

To determine mutation rates by amplicon sequencing, PCR was performed with target-specific primers (Table S1), amplifying approximately 225 bp around the cut site using Phusion High Fidelity (New England Biolabs, Ipswich, MA) polymerase. Primers contained a 5′-stub compatible with Illumina NGS library preparation. PCR products were ligated to Illumina TruSeq adaptors and purified. Libraries were prepared using a NEBNext kit (Illumina, San Diego, CA) according to the manufacturer’s guidelines. Samples were deep sequenced on an Illumina iSeq at 200 bp paired-end reads to a depth of approximately 10,000 reads per sample. Cortado (https://github.com/staciawyman/cortado) was used to analyze editing outcomes. Briefly, reads were adapter trimmed then merged using overlap to single reads. These joined reads were then aligned to the target reference sequence. Editing rates are calculated by counting any reads with an insertion or deletion overlapping the cut site or occurring within a 3 bp window on either side of the cut site. SNPs occurring within the window around the cut site are not counted. Total edited reads are then divided by the total number of aligned reads to derive percent edited.

### Western blot

Total plant tissue originating from 10 IEs at different timepoints were frozen in LN2, ground to a fine powder by mortar and pestle, and resuspended in 200 μl 2x Laemmli Sample Buffer (Bio-Rad, Hercules, CA) with 2-mercaptoethanol. Samples were boiled for 5 min, and the total soluble protein extracts (25 μl or 40 μl per well) were separated on 4-20% Mini-PROTEAN TGX precast polyacrylamide gels (Bio-Rad, Hercules, CA) and subsequently transferred to a 0.45 μm nitrocellulose membrane (GVS, Sanford, ME). For detection of Cas9 protein, anti-CRISPR/Cas9 C-terminal mouse monoclonal antibody (SAB4200751; Sigma-Aldrich, St. Louis, MO) and ProSignal Dura ECL Reagent (Genesee Scientific, San Diego, CA) were used. PageRuler Plus Prestained Protein Ladder (10–250 kDa, Thermo Fisher, Waltham, MA) was used as a molecular weight marker, and Cas9 protein with a C-terminal double nuclear-localization tag (QB3 Macrolab, University of California, Berkeley) was used as a positive control.

### Production of SnToxA

*SnToxA* was expressed in the *Pichia pastoris* yeast strain X33 (Liu et al. 2009) and cultured in yeast peptone dextrose broth (10 g yeast extract, 20 g peptone, 100 ml 20% dextrose in 900 ml distilled water) for 48 hours at 30 °C. Culture filtrate was harvested and filtered through a 0.45 µm HVLP filter membrane (Merk Millipore Ltd., Cork, Ireland) and dialyzed overnight against water using 3.5 kDa molecular weight cut off Snake Skin dialysis tubing (Thermo Scientific, IL, USA). Dialyzed filtrate was loaded onto a HiPrep SP XL 16/10 cation exchange column (GE Healthcare Piscataway, NJ). Unbound protein was washed off the column using a 20 mM sodium acetate (pH 5.0) buffer prior to a gradient elution of SnToxA using a buffer consisting of 300 mM sodium chloride and 20 mM sodium acetate (pH 5.0). Fractions that contained SnToxA were collected and frozen prior to lyophilizing to increase the concentration of SnToxA. Lyophilized samples were dissolved in a buffer consisting of 5 mM MOPS sodium salt (Alfa Aesar, MA, USA) and water, prior to infiltration into the plants.

### Production of SnTox5

*P. nodorum* strain Sn79+Tox5-3, generated by transforming *SnTox5* in to the avirulent *P. nodorum* strain Sn79-1087 (Kariyawasam et al. 2021),was used to prepare the culture filtrates containing SnTox5 as previously described (Friesen & Faris 2012) with minor modifications. In brief, Sn79+Tox5-3 was grown on V8-potato dextrose agar medium till spores were released from pycnidia. The plates were flooded with 10 ml of sterile distilled water, and 500 µl of spore suspension was used to inoculate 60 ml of liquid Fries medium (5 g ammonium tartrate, 1 g ammonium nitrate, 0.5 g magnesium sulfate, 1.3 g potassium phosphate [dibasic], 3.41 g potassium phosphate [monobasic], 30 g sucrose, 1 g yeast extract in 1000 ml of distilled water). Cultures were grown on an orbital shaker at 100 rpm for a week prior to two weeks of stationary growth under dark conditions at room temperature. Culture filtrates were filtered through a layer of Miracloth (EMD Millipore Corp, MA, USA) and were concentrated 5-fold using Amicon Ultracel – 3K centrifugal filters (Merk Millipore Ltd., Cork, Ireland). Culture filtrates were diluted in a 1:1 ratio with sterile water prior to infiltration into the plants.

### Necrotrophic effector infiltrations

Infiltrations with SnToxA and SnTox5 containing culture filtrates were conducted as previously described (Friesen & Faris 2012). Three infiltrations were performed per plant, and sensitivity was evaluated on a binary scale at 3 days post infiltration.

## Author Contributions

SP conceived and designed the experiments, analyzed the data, prepared the tables and figures, and wrote the manuscript with input from all co-authors. SP and JT performed the experiments. KLDR, GKK, JDF, TLF, and M-JC provided critical reagents, information, and discussion.

JHDC and BS supervised the work. All authors critically reviewed and edited the manuscript.

## Acknowledgements

We thank Innovative Genomics Institute (IGI) scientists, Netravathi Krishnappa and Stacia Wyman, for NGS library preparation and sequencing work and for assistance with analysis of sequencing data. We also thank Danielle J. Holmes from USDA-ARS for providing partially purified SnToxA. This work was supported by the IGI Founder’s Fund of the University of California at Berkeley.

## Supporting Information

**Table S1**

gRNA target sequences.

**Table S2**

Primers used to amplify the target region for amplicon next generation sequencing. Nucleotides shown in capital letters are the 5′-stub compatible with Illumina NGS library preparation.

**Table S3**

Genotypes of all edited M_0_ plants obtained. + indicates the number of base pairs inserted, - indicates the number of base pairs deleted.

**Figure S1**

Protoplast viability curve. N=3. Error bars indicate SEM.

**Figure S2**

Example of the difference in the number of unique mutant alleles between 14 dpb and 48 dpb. Provided are the detected alleles in immature embryos bombarded with Tsn1g2-Cas9 RNPs and treated at 37°C. The vertical bold dashed line represents the Cas9 cleavage site. Mutant alleles are marked with *. Wild type alleles are marked as WT. Dashes indicate base pair deletions, red boxes indicate base pair insertions, and bold letters indicate base pair substitutions.

