## Supplement for "Impact of temperature and time on DNA-free Cas9-ribonucleoprotein mediated gene editing in wheat protoplasts and immature embryos"

| gRNA ID | Target Site Sequence |
|---|---|
| Pi21gD | ACAACAGGGTGATCGTCCGT |
| Tsn1g2 | GGAACTGTCTACTAATATAT |
| Tsn1g3 | ACCATAAAGGGGATTTGTGA |
| Snn5g1 | ATACAATGGAGAACTAGTTA |
| Snn5g2 | CTTGCAAGGACTTGATGATA |

**Table S1**

| **Amplicon target** | **F Primer** | **R Primer** |
|---|---|---|
| Pi21gD | GCTCTTCCGATCTagttcttcttacgtaagattgatcata | GCTCTTCCGATCTcaggccttgcaccagatctt |
| Tsn1g2 / Tsn1g3 | GCTCTTCCGATCTggaaactgatttctc | GCTCTTCCGATCTcaaaatccgccagtt |
| Snn5g1 | GCTCTTCCGATCTtgacagtgaattccgtaacc | GCTCTTCCGATCTtagtaatgtggagcaccttc |
| Snn5g2 | GCTCTTCCGATCTgctgactacaaacagattgtcc | GCTCTTCCGATCTtaactatttggtagcagtagcc |

**Table S2**

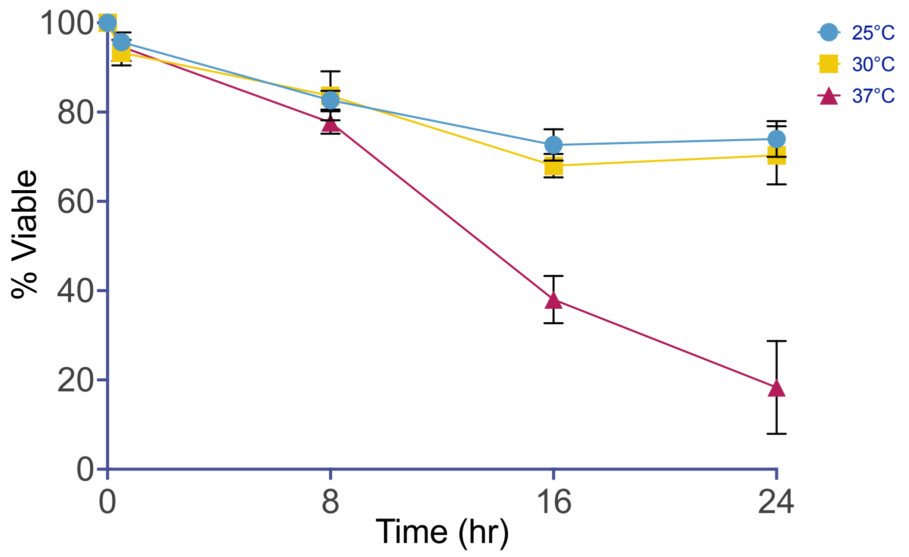


**Figure S1**

Tsn1g2 14 dpb IEs 37°C

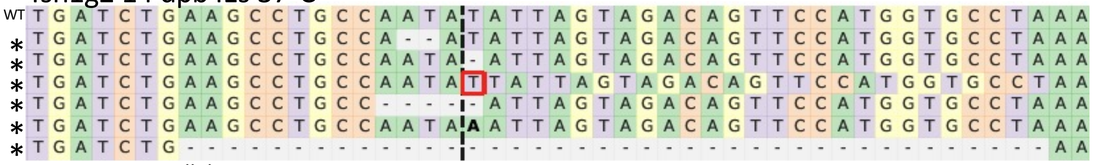


6 unique mutant alleles

Tsn1g2 48 dpb IEs 37°C

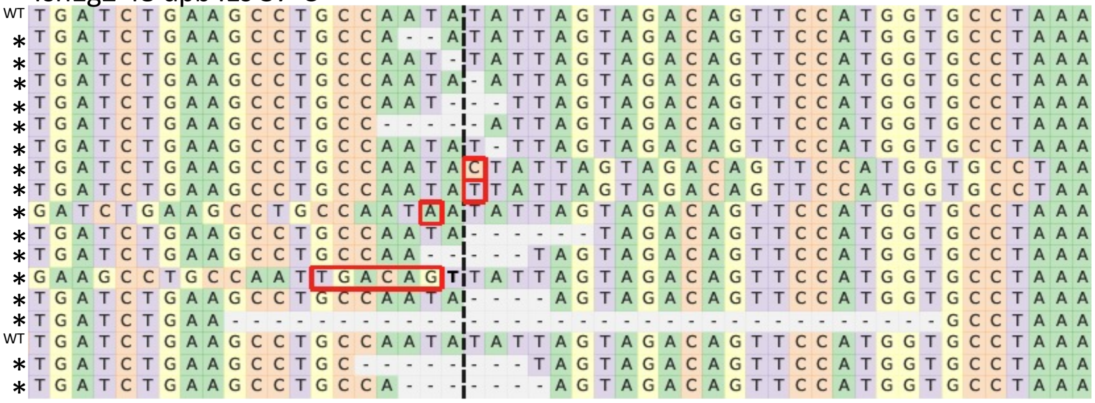


16 unique mutant alleles

**Figure S2**
